# Adolescent to young adult longitudinal development of subcortical volumes in two European sites with four waves

**DOI:** 10.1101/2021.06.09.447677

**Authors:** Lea L. Backhausen, Juliane H. Fröhner, Hervé Lemaître, Eric Artiges, Marie-Laure Palillère Martinot, Megan M. Herting, Fabio Sticca, Tobias Banaschewski, Gareth J. Barker, Arun L.W. Bokde, Sylvane Desrivières, Herta Flor, Antoine Grigis, Hugh Garavan, Penny Gowland, Andreas Heinz, Rüdiger Brühl, Frauke Nees, Dimitri Papadopoulos-Orfanos, Luise Poustka, Sarah Hohmann, Lauren Robinson, Henrik Walter, Jeanne Winterer, Robert Whelan, Gunter Schumann, Jean-Luc Martinot, Michael N. Smolka, Nora C. Vetter, the IMAGEN Consortium

**Author notes:** **Correspondence to** Lea L. Backhausen or Nora C. Vetter, Universitätsklinikum Carl Gustav Carus, Technische Universität Dresden, Fetscherstraße 74, 01307 Dresden, Germany, or. Authors contributed equally (Hervé Lemaître, Eric Artiges, Marie-Laure Palillère Martinot, Gareth J. Barker). Authors contributed equally (Jean-Luc Martinot, Michael N. Smolka).

## Abstract

Adolescent subcortical structural brain development might underlie psychopathological symptoms, which often emerge in adolescence. At the same time, sex differences exist in psychopathology, which might be mirrored in underlying sex differences in structural development. However, previous studies showed inconsistencies in subcortical trajectories and potential sex differences. Therefore, we aimed to investigate the subcortical structural trajectories and their sex differences across adolescence using for the first time a single cohort design, the same quality control procedure, software and a general additive mixed modeling approach. We investigated two large European sites from ages 14 to 24 with 503 participants and 1408 total scans from France and Germany as part of the IMAGEN project including four waves of data acquisition. We found significantly larger volumes in males versus females in both sites and across all seven subcortical regions. Sex differences in age-related trajectories were observed across all regions in both sites. Our findings provide further evidence of sex differences in longitudinal adolescent brain development of subcortical regions and thus might eventually support the relationship of underlying brain development and different adolescent psychopathology in boys and girls.

## 1 Introduction

Adolescence is a core incidence phase for the development of mental disorders (Paus et al., 2008). Mental disorders occur in around 20% of adolescents with important sex differences, i.e. higher rates of internalizing disorders in girls and externalizing disorders in boys (Kessler et al., 2005; Zahn-Waxler et al., 2008). These sex differences might be mirrored in underlying sex differences in subcortical structural development (Gogtay & Thompson, 2010; Paus et al., 2008; Shaw et al., 2010) since alterations in subcortical structures have been associated with psychopathological disorders (Hoogman et al., 2017; Noordermeer et al., 2017; Rogers & Brito, 2016). Hence, there is a need to characterize typical subcortical structural development and the related sex differences across adolescence to better understand adolescent psychopathology.

Cross-sectional (Boedhoe et al., 2020; Brain Development Cooperative Group, 2012; Østby et al., 2009) and longitudinal studies (Dennison et al., 2013; Goddings et al., 2014; Herting et al., 2018; Narvacan et al., 2017; Tamnes et al., 2013; Wierenga et al., 2014) began to investigate the typical development of subcortical structures across adolescence with some studies also analyzing sex effects (Goddings et al., 2014; Herting et al., 2018; Narvacan et al., 2017; Wierenga, Bos, et al., 2018; Wierenga et al., 2014). Differential trajectories across adolescence have been revealed for some subcortical structures. Importantly, we review here the trajectories from the age span 14 to 24, which will also be covered by our study.

The basal ganglia (mainly caudate nucleus, putamen, globus pallidus, and nucleus accumbens) showed volume decreases across adolescence in cross-sectional studies (Brain Development Cooperative Group, 2012; Østby et al., 2009), including a life span mega-analysis (Dima et al., 2021). A longitudinal study with 85 participants from 8-22 years and two waves (170 scans) corroborated these findings (Tamnes et al., 2013). Other longitudinal studies showed similar decreases across adolescence for some basal ganglia structures (Dennison et al., 2013: 120 scans, two waves, n=60, 11-17 years; Goddings et al., 2014: 711 scans, ≥ two waves, n= 275, 7-19 years; Narvacan et al., 2017: 168 scans, two waves, n=84, 5-27 years; Wierenga et al., 2014: 223 scans, up to three waves, n=147, 7–24 years; Raznahan et al., 2014: decrease for the globus pallidus only: 1172 scans, n=618, 5–25 years, about half having ≥ two waves; Wierenga et al., 2018: decrease for the accumbens only: 680 scans, n= 271, 8–28 years, up to three scans).

However, diverging findings from the overall decreasing pattern exist. The caudate nucleus remained stable across adolescence in three studies: for both sexes (Narvacan et al., 2017; Wierenga, Bos, et al., 2018) or for males only (Raznahan et al., 2014). For the globus pallidus three studies found volume increase (Dennison et al., 2013; Wierenga, Bos, et al., 2018; Wierenga et al., 2014) with Wierenga et al. (2014) showing a peak at age 17 with a slight decrease thereafter. A multi-site study found a descriptive, albeit non-significant increase (Herting et al., 2018: three sites, n=216 participants and 467 scans, 10-22 years, two to three waves). For the nucleus accumbens a stable trajectory was found for both sexes (Dennison et al., 2013) or for males only (Herting et al., 2018). For the putamen a non-linear increase was found (Wierenga, Bos, et al., 2018).

For the thalamus, previous results converged in indicating volume decreases from about age 13 (Tamnes et al., 2013) or age 15 to 19 on (Herting et al., 2018 for males only; Raznahan et al., 2014; Wierenga, Bos, et al., 2018) while Narvacan et al. (2017) only showed decreases in their cross-sectional cohort and no changes in their longitudinal cohort.

Diverging findings also exist regarding the trajectories of the amygdala and hippocampus across adolescence (Koolschijn & Crone, 2013; Østby et al., 2009; Sowell et al., 2002; Wierenga et al., 2014). While some cross-sectional (Durston et al., 2001; Giedd et al., 1996) and longitudinal studies (Dennison et al., 2013; Herting et al., 2018) found an increase of the amygdala or hippocampus during adolescence, other longitudinal studies showed decreases (Tamnes et al., 2013) or little to no change (Dennison et al., 2013; Wierenga, Bos, et al., 2018). For both the amygdala and hippocampus a slight decrease for girls at about 15 years versus an increase for boys has been shown in another study (Goddings et al., 2014). Wierenga et al. (2014) reported an increase for both the amygdala and hippocampus with a peak at about age 17/ 18 followed by a slight decrease.

### 1.1 Sex differences

Regarding sex differences, cross-sectional and longitudinal studies consistently demonstrate larger subcortical volumes in boys versus girls across adolescence when not correcting for whole brain size (Dennison et al., 2013; Goddings et al., 2014; Herting et al., 2018; Narvacan et al., 2017; Raznahan et al., 2014; Tamnes et al., 2013; Wierenga, Bos, et al., 2018; Wierenga et al., 2014) ranging from 9% for the putamen to 15% for the amygdala (Narvacan et al., 2017).

There has also been note of sex differences in age-related trajectories of subcortical structures; however, previous studies are rather inconsistent. Overall differences in trajectories between sexes were found in some studies (Dennison et al., 2013; Goddings et al., 2014; Herting et al., 2018) and others reported later peak volumes in males (Lenroot et al., 2007; Raznahan et al., 2014). The thalamus declined for females (Dennison et al., 2013) or males only (Herting et al., 2018) when tested separately for the sexes. A slight decrease for the amygdala and hippocampus was found in girls versus a continued increase for boys (Goddings et al., 2014). Significantly differing nonlinear trajectories across sexes were also found for the amygdala and hippocampus (Herting et al., 2018). Further, no sex differences in trajectories have also been reported (Narvacan et al., 2017; Wierenga, Bos, et al., 2018; Wierenga et al., 2014).

Importantly, in most studies the sex differences in trajectories can only be inferred indirectly when descriptively comparing shapes of trajectories. Only very few studies tested for statistical differences in trajectories between the sexes (Herting et al., 2018; Wierenga, Bos, et al., 2018). Most of the studies further had no or a very low amount of participants with more than two waves (except Goddings et al., 2014; Raznahan et al., 2014; Wierenga, Sexton, et al., 2018), limiting their conclusions on trajectories and their ability to model more complex trajectories. Moreover, some studies only covered few subcortical regions (four or five: Goddings et al., 2014; Raznahan et al., 2014), which limits inferences about trajectories of all subcortical structures.

Taken together, previous studies showed inconsistent results in subcortical age trajectories and their potential sex differences. This might be related to differences in sample characteristics, study design or analysis regarding sampling type, age and sex, sociodemography, inter-scan interval, number of total scans, image acquisition, quality control, segmentation software, and analysis models. Regarding sampling type, most studies used accelerated longitudinal designs (Herting et al., 2018; Narvacan et al., 2017; Raznahan et al., 2014; Wierenga, Bos, et al., 2018). These designs are limited because age effects are confounded with interactions between a given starting age and when the measurement happened and participants only contribute data to parts of the time span investigated. Some studies included differing sex ratios across ages (Herting et al., 2018) or did not report detailed sociodemographic sample description like IQ or socioeconomic status (Herting et al., 2018; Tamnes 2013) limiting understanding of the generalizability of findings. Image acquisition differed in field strengths or scanner types with most studies using 1.5 Tesla (except Wierenga et al. (2018) using 3 Tesla) and some studies using different scanners within studies (Dennison et al., 2013, using two different 3 Tesla scanners and Herting et al. (2018) with two sub-sites using two different 3 Tesla scanners; see Liu et al., 2020 for the influence of scanner type differences). Differences in quality control procedures may have also impacted the results (Backhausen et al., 2016; Ducharme et al., 2016). The type of segmentation software can also influence volume estimates (Makowski et al., 2018; Morey et al., 2010), even across different versions of the same software (e.g. differences in putamen using FreeSurfer 5.3 versus 6.0, see release notes). Only in some analysis software packages a longitudinal processing approach is included (see Vijayakumar et al., 2018 for a software overview) and thus has not always been used. Previous studies further employed different analysis models, e.g. linear mixed effects modelling (Goddings et al., 2014; Narvacan et al., 2017) or hierarchical regression (Dennison et al. 2013). These different analysis approaches may impact results (Vijayakumar et al., 2018). Parametric models use the a priori assumption of either linear, quadratic, or cubic age-related change and assume similar patterns of age trajectories between groups. In contrast, generalized additive mixed models (GAMMs) are more flexible since they fit curves using ‘smooth function’ terms and thus have recently become more widely applied (e.g. Herting et al., 2018; Wierenga, Bos, et al., 2018; Wierenga et al., 2014). GAMMs describe the best relationship between predictor and outcome variables of interest without a priori knowledge of the inherent form of the data. Therefore, GAMMs allow for different nonlinear shapes of age trajectories for categories of a variable such as sex with the categories males and females.

Taken together, the methodical differences between studies still limit our ability to draw conclusions about the subcortical structural trajectories in the mid-adolescent to young adult age span, specifically due to the lower amount of studies with more than two waves including a meaningful amount of participants in this age span. Hence, there is still the need to test for possible sex differences in subcortical age trajectories across adolescence. Therefore, we aimed to investigate the subcortical structural trajectories including the thalamus, globus pallidus, caudate nucleus, putamen, nucleus accumbens, hippocampus, and amygdala as regions of interest (ROIs) and their sex differences across adolescence using the same quality control procedure, segmentation software, i.e. FreeSurfer 6.0.0, and a GAMM approach in a unique study of adolescents from two European sites with large samples. Importantly, differing from most previous accelerated studies, our design was a single cohort approach across two sites where all participants started at the same age and were followed across the entire age-range of interest. Specifically, as part of the Imaging Genetics (IMAGEN) project (Schumann et al., 2010) data stemmed from a site in Paris, France (N=255) and a site in Dresden, Germany (N=248) including four waves of data acquisition at the ages of M=14.47 years (range 13.33 – 15.72), M=16.48 years (range 15.65 – 17.88), M=19.24 years (range 17.26 – 22.54) and M=22.35 years (range 20.1 – 24.76) across sites, with a total of 1408 scans. The sites were largely homogeneous in terms of age range, sex distribution, inter-scan interval, number of total scans and sociodemographic characteristics such as socioeconomic status and ethnicity. In examining two similar sites, we can extend previous research that showed sex differences in trajectories dependent on specific sites (Herting et al., 2018) and thus explore age-constant and time-varying site differences closely focused on the mid-adolescent to young adult age span with more waves and scans within this particular age-range enabling more precise estimation of trajectories.

## 2 Methods

Access to the IMAGEN dataset is available with an accepted proposal from the IMAGEN executive committee (https://imagen-europe.com/resources/imagen-project-proposal/). All code is posted to Open Science Framework and can be assessed at https://osf.io/5tfh4/.

### 2.1 Participants and study design

The study is part of the larger European IMAGEN project assessing magnetic resonance imaging (MRI) data from typically developing adolescents at the age of 14, 18, and 22. For a detailed description of recruitment and assessment procedures in the IMAGEN study, please refer to (Schumann et al., 2010). At the sites Dresden and Paris MRI data was assessed additionally at age 16, resulting in four waves (see Table 1 for sample characteristics and Figure 1 for age and sex distributions per site), therefore these two sites were chosen. Participants provided written informed consent and assent (in case of participants <18 years also their legal guardians). The study had been approved by the local ethics committees (Technische Universität Dresden and University of Paris) and was performed in accordance with the Declaration of Helsinki. Exclusion criteria included existing bipolar disorder, schizophrenia and major neuro-developmental disorders such as autism, as well as a premature birth, head trauma and history of several neurological or medical disorders. In this manuscript, female and male is defined as “assigned female at birth” and “assigned male at birth”, respectively. For Dresden, 260 participants took part in the overall IMAGEN study of whom 256 were scanned for at least one wave and 248 participants were included in the analyses after quality control. For Paris, 274 participants took part in the study of whom 258 were scanned for at least one wave and 255 participants were included in the analyses after quality control (see below and supplementary Table S1 for details). In total, the present study included 503 participants (250 females) and 1408 scans covering the age range of 13.33–24.76 years, including 447 scans at wave 1 (Dresden: 201, Paris: 246), 336 scans at wave 2 (Dresden: 207, Paris: 129), 372 scans at wave 3 (Dresden: 172, Paris: 200), and 253 scans at wave 4 (Dresden: 130, Paris: 123). Of the 503 participants included in the study 94 % were White (97% Dresden, 90% Paris) and the majority stemmed from rather well-educated households (as a proxy for socioeconomic status), with around 60% of the parents having obtained a university or college (university of applied sciences) degree. For more information about the parental education assessment in the IMAGEN study see Table 1 in Schumann et al. (2010).

**Table 1.** Sample characteristics. Sample characteristics including demographics of typically developing adolescents from two sites of the IMAGEN study using single cohort designs.

|  | Dresden, Germany |  |  | Paris, France |  |  |
| --- | --- | --- | --- | --- | --- | --- |
|  | All | Female | Male | All | Female | Male |
| Total scans across all waves | 710 | 352 | 358 | 698 | 366 | 332 |
| Age (years) |  |  |  |  |  |  |
| Wave 1, M (SD) | 14.54 (.32) | 14.58 (.32) | 14.49 (.33) | 14.42 (.5) | 14.43 (.5) | 14.41 (.5) |
| Wave 2, M (SD) | 16.6 (.42) | 16.64 (.4) | 16.56 (.42) | 16.80 (.62) | 16.84 (.59) | 16.76 (.64) |
| Wave 3, M (SD) | 18.68 (.57) | 18.72 (.56) | 18.64 (.58) | 19.73 (.73) | 19.76 (.74) | 19.69 (.73) |
| Wave 4, M (SD) | 22.1 (.74) | 22.04 (.62) | 22.12 (.85) | 22.64 (.55) | 22.62 (.52) | 22.67 (.59) |
| N |  |  |  |  |  |  |
| Total | 248 | 121 | 127 | 255 | 129 | 126 |
| Participants with 1 scan | 35 | 14 | 21 | 41 | 17 | 24 |
| Participants with 2 scans | 47 | 26 | 21 | 51 | 25 | 28 |
| Participants with 3 scans | 83 | 38 | 45 | 95 | 49 | 44 |
| Participants with 4 scans | 83 | 43 | 40 | 68 | 38 | 30 |
| Demographics - *Data from the first wave is reported\** |  |  |  |  |  |  |
| Pubertal status<sup>a</sup>, M(SD) | 3.65 (.66) | 4.03 (.41) | 3.28 (.64) | 3.55 (.78) | 3.98 (.42) | 3.1 (.82) |
| Height in cm, M(SD) | 169.42 (7.21) | 166.55 (6.45) | 172.13 (6.85) | 165.57 (8.03) | 162.67 (5.54) | 168.6 (9.07) |
| Weight in kg, M(SD) | 59.17 (11.74) | 56.82 (10.23) | 61.39 (12.64) | 55.3 (9.26) | 53.61 (7.9) | 57.05 (10.25) |
| BMI, M(SD) | 20.53 (3.29) | 20.42 (3.03) | 20.62 (3.51) | 20.11 (2.61) | 20.23 (2.6) | 19.98 (2.65) |
| IQ<sup>b</sup>, M(SD) | 112.95 (10.72) | 113.05 (9.93) | 112.86 (11.48) | 109.35 (9.43) | 108.5 (8.95) | 110.25 (9.87) |

\*Note that data for pubertal status, height, weight, BMI and IQ is reported for all participant in the final data set independent of a high-quality scan at the first wave. BMI = Body mass index (BMI = weight in kg/ height in m<sup>2</sup>).
<sup>a</sup>Pubertal status ranged from 1 for prepubertal to 5 for postpubertal status, measured with the Pubertal Development Scale (PDS; Petersen et al., 1988)
<sup>b</sup>General cognitive ability estimated with the subtests Similarities, Block Design, Vocabulary, and Matrices from the Wechsler Intelligence Scale For Children (WISC-IV, German adaptation; Petermann & Petermann, 2007)
\* $p < .05$ , \*\* $p < .01$ , \*\*\* $p < .001$ .

**Figure 1:**
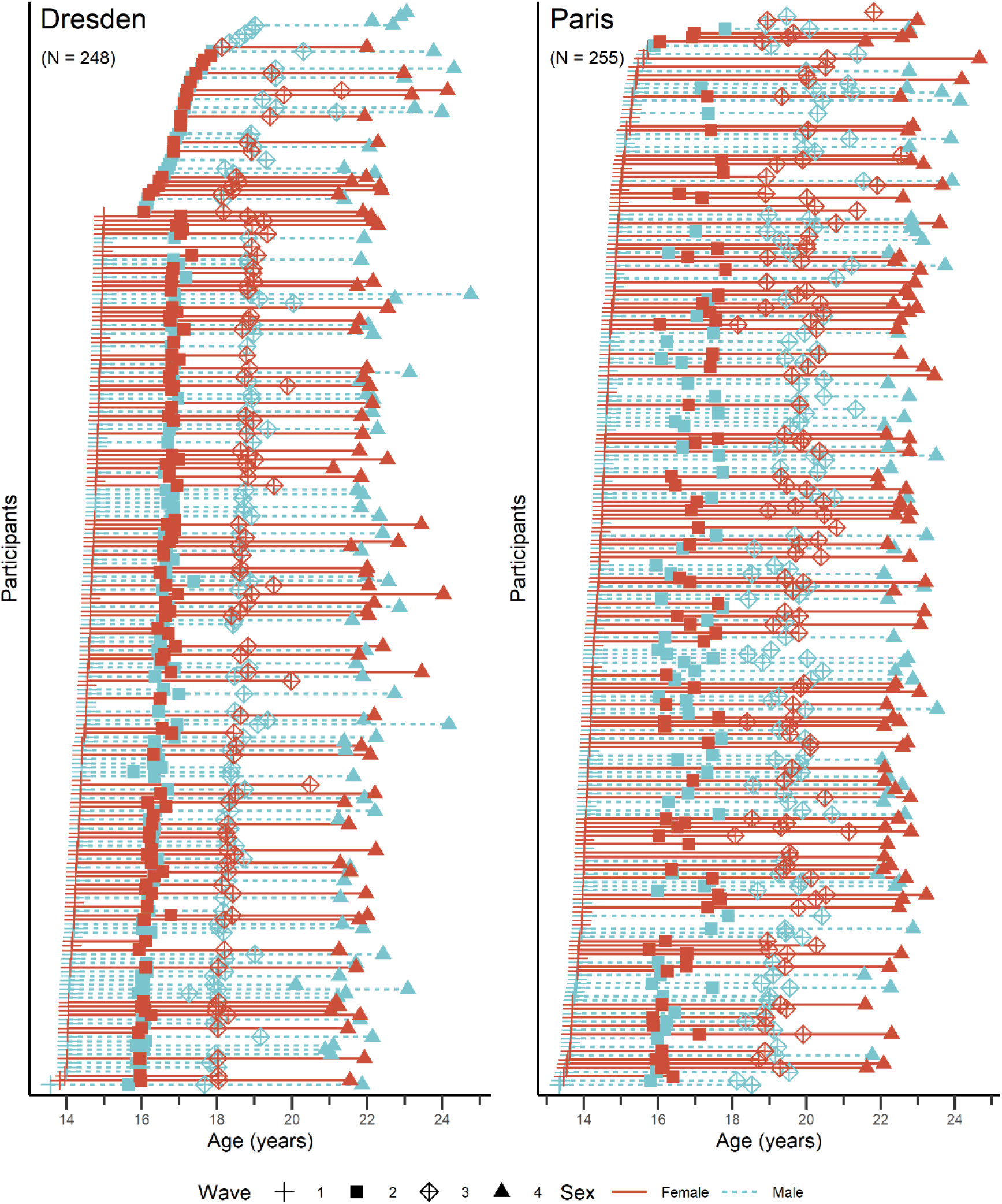
Age and sex distributions for each site.

### 2.2 Structural imaging

#### 2.2.1 Image acquisition

We acquired T1-weighted anatomical scans using the respective local Siemens (Erlangen, Germany) 3T whole-body MR tomograph equipped with a 12-channel head coil in Dresden and a 64-channel head coil in Paris. While in Dresden this was a MAGNETOM Trio for all waves, the MAGNETOM Trio was updated to a MAGNETOM Prisma during wave 3. For both sites a standardized quality control assessment for the scanner was regularly performed (phantom scan and in-vivo scan, see Schumann et al. (2010)). High-resolution T1-weighted images were collected using a magnetization prepared rapid acquisition gradient-echo (MPRAGE) sequence [Dresden/ Paris: repetition time (TR) = 1900/ 2300 ms, echo time (TE) = 2.26/ 2.93 ms, inversion time (TI) = 900/ 900 ms, voxel size = 0.5 x 0.5 x 1.0/ 1.1 x 1.1 x 1.1 mm, flip angle = 9/ 9°; matrix size = 256 x 256/ 256 x 256 mm; 176/ 160 slices]. The sequence in Dresden varied slightly. This is due to differences in imaging data assessment waves at the each site of the IMAGEN project. Only the sites Dresden and Paris assessed imaging data at age 16 with Dresden using the local MPRAGE sequence only. All images were examined by a clinical neuroradiologist for structural abnormalities.

#### 2.2.2 Image processing

Images (also single time points) were processed using the longitudinal pipeline of FreeSurfer 6.0.0 (http://surfer.nmr.mgh.harvard.edu; Fischl et al., 2002; Reuter et al., 2012). This pipeline creates an unbiased within-subject template space and image using robust, inverse consistent registration (Reuter et al., 2010). Several processing steps, such as skull stripping, Talairach transforms, atlas registration, as well as spherical surface maps and parcellations are then initialized with common information from the within-subject template. This process significantly improves robustness with respect to noise, intensity scaling and outliers, significantly increasing statistical power (Reuter et al., 2012) and scan-rescan reliability as compared to conventional cross-sectional analysis pipelines (Jovicich et al., 2013). We further used the recon-all-flag-3T which alters FreeSurfer’s internal N3 bias field correction parameters (Sled et al., 1998) making it more appropriate for 3T MRI (Zheng et al., 2009).

As previous studies demonstrated no evidence of developmental difference between hemispheres in our subcortical ROIs (Brain Development Cooperative Group, 2012; Østby et al., 2009), we report averaged volumes over both hemispheres similarly to previous studies (Herting et al., 2018; Wierenga, Bos, et al., 2018). Correlations between hemispheres across sites at each wave were high for each ROI (*r’s* > .7) except for the nucleus accumbens (*r’s* = .46 to .79) and the amygdala (*r’s* = .6 to .79; see supplementary Table S2 for all correlations). Following the methodology of previous studies (e.g. Herting et al., 2018; Wierenga, Bos, et al., 2018) we used absolute volume values and did not correct ROI volumes for intra-cranial volume.

#### 2.2.3 Quality control

Before statistical analyses, trained operators performed the following pre- and post-processing quality control: For pre-processing quality control all raw images were visually inspected for motion and technical artifacts according to our lab’s three-category (fail, check, pass) rating scheme (Backhausen et al., 2016). We retained “check” and “pass” images and excluded “fail” images from longitudinal pipeline processing to ensure high quality of the within-subject template. In post-processing quality control trained operators inspected longitudinally processed images with questionable raw image quality (rated “check”) for accuracy of subcortical segmentation. Subcortical structures were excluded when incorrectly segmented (see supplementary Table S1). No manual adjustments of subcortical volumes were made.

#### 2.2.4 Statistical analyses

For analysis of GAMMs we used R version 3.6.1 with the mgcv package version 1.8-33 (Wood, 2004, 2006) and the packages itsadug version 2.3 and ggplot2 version 3.3.2 for visualization of smooth curves. We used a shrinkage version of penalized cubic regression splines as smooth terms in all models. The k parameters, which specify the number of basis functions to build the curves and thus influence potential over-fitting, were set to 4 after k.check (mgcv package) analyses.

We closely followed the approach of Herting et al. (2018) and built GAMM models to examine age-constant and time-varying sex and site differences in trajectories of subcortical volumes. We explored differences in trajectories by examining significance of difference smooths between the trajectories of males versus females in the Dresden and Paris site. GAMM models were implemented to (1) test an age-constant sex difference (main effect of sex) as well as a sex difference in the age trajectory (age*sex interaction), while controlling for site at the level of main effect and site*interaction and (2) test an age-constant site difference (main effect of site) as well as a site difference in the age trajectory (age*site interaction), while controlling for sex at the level of main effect and sex*age interaction. All models included an individual-level random effect intercept per participant. Sex and site were coded as factors (male = 0, female = 1; Dresden = 0, Paris = 1). To directly test sex and site differences, previous GAMM models were updated in coding sex/ site and sex*age /site*age as contrasting ordered factors. For GAMM estimates of these smooth terms see Table 2.

**Table 2.** GAMMS examining sex and age effects across both sites on subcortical regions of interest.

| THALAMUS |  |  |  |  |
| --- | --- | --- | --- | --- |
| Main effect | $\beta$ | SE | $t$ | $p$ |
| Sex (F vs M) | -734.91 | 53.75 | -13.67 | *** |
| Trajectory | edf | Ref.df | $F$ | $p$ |
| S(age) | 2.85 | 3 | 24.44 | *** |
| S(age, F vs M) | 2.29 | 3 | 11.44 | *** |
| GLOBUS PALLIDUS |  |  |  |  |
| Main effect | $\beta$ | SE | $t$ | $p$ |
| Sex (F vs M) | -184.97 | 17.39 | -10.64 | *** |
| Trajectory | edf | Ref.df | $F$ | $p$ |
| S(age) | 0.81 | 3 | 0.45 | ** |
| S(age, F vs M) | 2.46 | 3 | 56.84 | *** |
| CAUDATE NUCLEUS |  |  |  |  |
| Main effect | $\beta$ | SE | $t$ | $p$ |
| Sex (F vs M) | -302.60 | 40.35 | -7.50 | *** |
| Trajectory | edf | Ref.df | $F$ | $p$ |
| S(age) | 1.88 | 3 | 1.78 | *** |
| S(age, F vs M) | 2.62 | 3 | 40.02 | *** |
| PUTAMEN |  |  |  |  |
| Main effect | $\beta$ | SE | $t$ | $p$ |
| Sex (F vs M) | -595.96 | 45.89 | -12.99 | *** |
| Trajectory | edf | Ref.df | $F$ | $p$ |
| S(age) | 1.76 | 3 | 1.53 | *** |
| S(age, F vs M) | 2.53 | 3 | 20.27 | *** |
| NUCLEUS ACCUMBENS |  |  |  |  |
| Main effect | $\beta$ | SE | $t$ | $p$ |
| Sex (F vs M) | -63.17 | 9.0 | -7.02 | *** |
| Trajectory | edf | Ref.df | $F$ | $p$ |
| S(age) | 0.00 | 3 | 0.00 | 0.62 |
| S(age, F vs M) | 1.80 | 3 | 11.52 | *** |
| HIPPOCAMPUS |  |  |  |  |
| Main effect | $\beta$ | SE | $t$ | $p$ |
| Sex (F vs M) | -360.67 | 30.39 | -11.87 | *** |
| Trajectory | edf | Ref.df | $F$ | $p$ |
| S(age) | 2.31 | 3 | 1.88 | 0.55 |
| S(age, F vs M) | 2.33 | 3 | 19.17 | *** |
| AMYGDALA |  |  |  |  |

| **Main effect** | $\beta$ | SE | $t$ | $p$ |
| --- | --- | --- | --- | --- |
| Sex (F vs M) | -238.48 | 15.28 | -15.61 | *** |
| **Trajectory** | edf | Ref.df | $F$ | $p$ |
| S(age) | 0.00 | 3 | 0.00 | 0.76 |
| S(age, F vs M) | 2.27 | 3 | 23.22 | *** |

GAMM estimates for age, sex, and age\*sex across both sites. Sex was coded as a factor (male = 0, female =1), allowing for each term to reflect the following: Sex (FvsM) reflected the main difference in females as compared to males; S(age) reflected the global smooth trajectory of age; S(age, F vs M) reflected the difference in trajectory of females compared males. Site and site\*age were included as covariates in all models. Estimate, standard error (SE), $t$ -value and associated $p$ -values for each main difference. Smooth function (S(age) / edf), degrees of freedom (Ref.df), $F$ -statistic and associated $p$ -values for each smooth term. \* $p < .05$ , \*\* $p < .01$ , \*\*\* $p < .001$ .

To test whether the inclusion of age*sex interaction terms significantly improve model fit and explain variance, we analyzed three models following procedures by Pedersen et al. (2019) and Wierenga et al. (2018; 2019) for each subcortical structure as a smooth function of age of the individual *i* at wave *j*, a random intercept per participant effect (*u_i_*) and error (*error_ij_*), while controlling for site (in main effect and site*age interaction) in each model. We compared these models using Akaike Information Criterion (AIC; Akaike, 1973). The model that had the lowest AIC score and significantly differed from more parsimonious models was selected as the best fit model.

Model 1: age only model including only a global smooth term for age

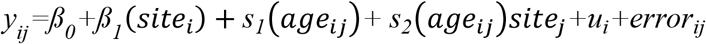

Model 2: age plus main effect of sex model including a global smooth term for age and a main effect of sex

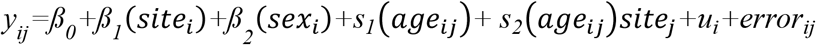

Model 3: age plus sex*age interaction including a global smooth term for age, a main effect of sex, and group-level smooth terms for males and females with differing complexity

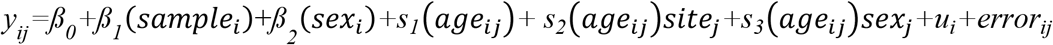

Here, *s_1_*, *s_2_*, and *s_3_* denote the arbitrary smooth functions (shrinkage version of cubic regression splines). Further, *β_0_* stands for the random intercepts per participant, *β_1_* for parameter estimates of site, and *β_2_* for parameter estimates of sex.

## 3 Results

For all seven subcortical structures, models with the interaction age*sex (Model 3) had lower AIC values than the more parsimonious models, and were thus deemed best fitting models (supplementary Table S3). Moreover, for all seven ROIs we found significant sex differences in age trajectories when examining difference smooths between females and males. Hence, we present results for GAMM models including age, sex, as well as an age*sex interaction to investigate age-constant sex differences as well as a sex difference in age trajectories (age*sex interaction), while controlling for site (main effect and age*site interaction). Notably, trajectories for each sex follow the global age trajectory (i.e. across all participants of both sexes) as they are penalized by the model to not differ too much from it (Pedersen et al., 2019). To better understand these differences, we examined age trajectories in GAMMs for each sex separately (independent from each other and the global age trajectory) while controlling for site (see Table 3 and Figure 2).

**Table 3.** GAMM estimates for age across both sites in females and males separately.

| FEMALES |  |  |  |  | MALES |  |  |  |  |
| --- | --- | --- | --- | --- | --- | --- | --- | --- | --- |
| THALAMUS | edf | Ref.df | *F* | *p* |  | edf | Ref.df | *F* | *p* |
| S(age) | 2.77 | 3 | 54.82 | *** | S(age) | 2.58 | 3 | 5.18 | *** |
| GLOBUS PALLIDUS |  |  |  |  |  |  |  |  |  |
| S(age) | 2.06 | 3 | 58.88 | *** | S(age) | 2.70 | 3 | 80.86 | *** |
| CAUDATE NUCLEUS |  |  |  |  |  |  |  |  |  |
| S(age) | 2.70 | 3 | 197.81 | *** | S(age) | 2.75 | 3 | 177.71 | *** |
| PUTAMEN |  |  |  |  |  |  |  |  |  |
| S(age) | 2.58 | 3 | 127.74 | *** | S(age) | 2.68 | 3 | 61.43 | *** |
| NUCLEUS ACCUMBENS |  |  |  |  |  |  |  |  |  |
| S(age) | 1.75 | 3 | 10.61 | *** | S(age) | 0.00 | 3 | 0.00 | 0.78 |
| HIPPOCAMPUS |  |  |  |  |  |  |  |  |  |
| S(age) | 2.45 | 3 | 4.22 | ** | S(age) | 1.76 | 3 | 1.15 | *** |
| AMYGDALA |  |  |  |  |  |  |  |  |  |
| S(age) | 0.00 | 3 | 0.00 | 0.74 | S(age) | 2.70 | 3 | 18.82 | *** |

Smooth function (S(age) / edf), degrees of freedom (Ref.df), F-statistic and associated p-values for each term. \* $p < .05$ , \*\* $p < .01$ , \*\*\* $p < .001$ .

**Figure 2.**
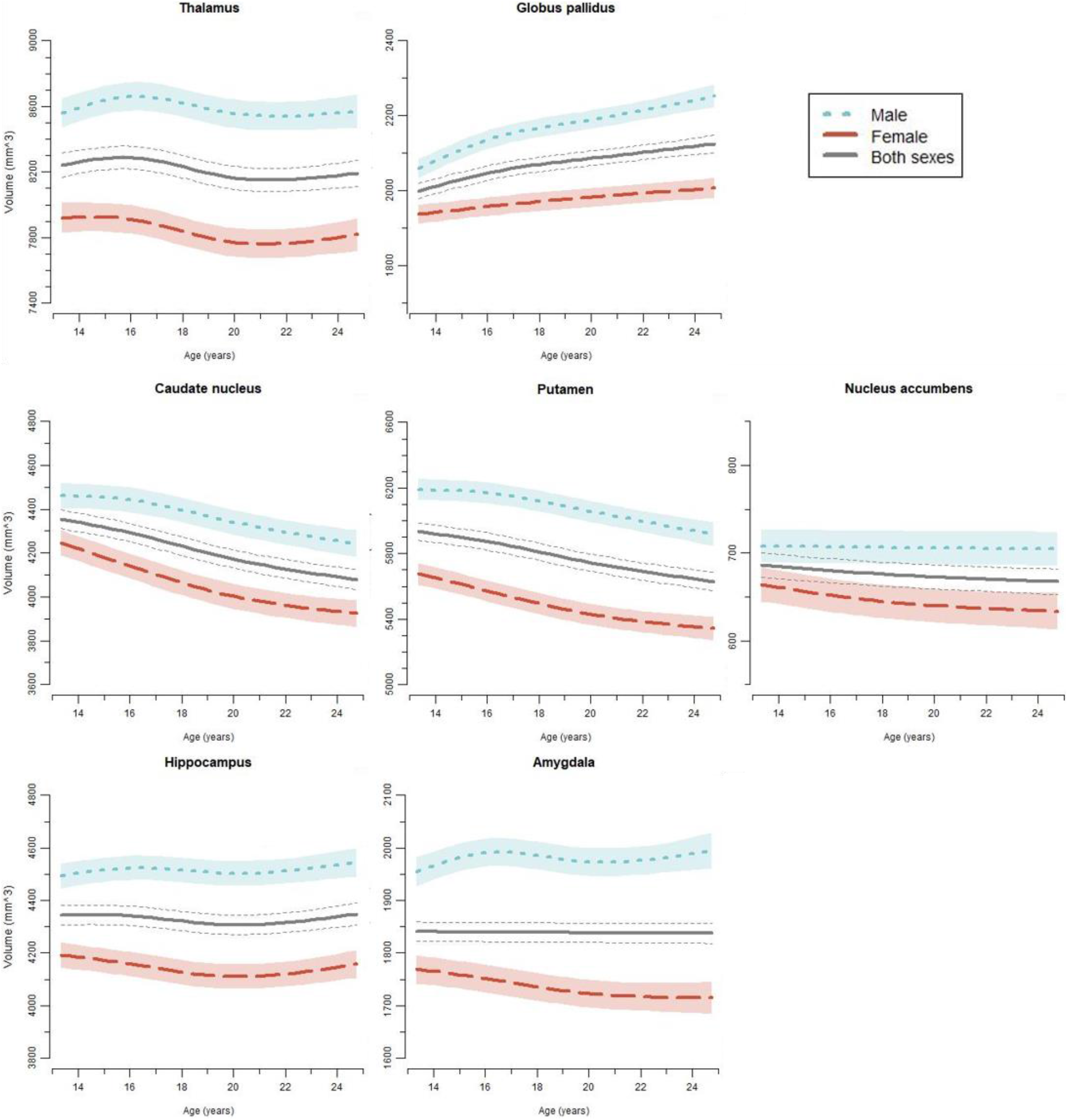
Age trajectories for subcortical ROIs across the two sites Dresden and Paris. GAMM fitting with 95% confidence intervals plotted separately for females, males, and both sexes across Dresden and Paris.

### 3.1 Sex differences in raw volumes

To investigate volume differences between the sexes for all data sets and in each wave separately, we calculated percentage differences between males and females from raw values, not from GAMM modeling. For all structures, males had larger volumes as compared to females across the entire age range with overall differences ranging from 8.7% for the hippocampus to 13.7% for the amygdala (see Table 4). Percentage differences increased with age in thalamus, globus pallidus, caudate nucleus, hippocampus, and amygdala, which is partly mirrored by increasingly diverging trajectories of females and males over time (see Table 4; Figure 2).

**Table 4.** Percentage volume differences of subcortical brain structures between sexes, ordered by largest to smallest in the overall difference.

| Structure | Difference in volume (male – female, %) |  |  |  |  |
| --- | --- | --- | --- | --- | --- |
|  | All<br>(N = xx) | TP1<br>(N = xx) | TP2<br>(N = xx) | TP3<br>(N = xx) | TP4<br>(N = xx) |
| Amygdala | 13.7 | 12.3 | 12.4 | 15.4 | 15.2 |
| Putamen | 10.8 | 9.4 | 10.6 | 12.5 | 10.7 |
| Thalamus | 9.8 | 8.6 | 9.7 | 10.2 | 11.6 |
| Pallidum | 9.4 | 7.2 | 9.0 | 11.0 | 11.8 |
| Nucleus accumbens | 9.2 | 8.0 | 8.8 | 11.9 | 7.9 |
| Hippocampus | 8.8 | 7.7 | 8.2 | 9.6 | 10.2 |
| Caudate nucleus | 7.4 | 5.6 | 7.5 | 8.7 | 8.2 |

For n of each structure per wave please refer to supplementary Table S1.

### 3.2 Trajectories for all participants; i.e. not differentiating sex and site

Estimation of smooth terms including all participants indicated overall volume decrease in caudate nucleus and putamen and overall volume increase in globus pallidus. Further, we found a nonlinear pattern of change in the thalamus and no significant volume change in nucleus accumbens, hippocampus and amygdala.

### 3.3 Sex differences in age trajectories

#### Thalamus

Males showed a volume increase until about age 17, while females remained rather stable until about age 16. After this, both sexes followed a similar nonlinear trajectory starting with a decrease and then a slight increase.

#### Globus pallidus

While females showed a nonlinear increase across the entire age range, the curve for males increased steeper until age 16 and flattened afterwards.

#### Caudate nucleus

For females, volume decreased until about age 22, then decrease flattened. Males showed a flatter decrease until about age 17, followed by a nonlinear decrease.

#### Putamen

For females, volume decreased until about age 20, then decrease flattened. Males showed a flatter decrease until about age 16, followed by a nonlinear decrease.

#### Nucleus accumbens

We did not find volume change in males, but a significant volume decrease in females which differed from the global age trajectory.

#### Hippocampus

While both groups showed significant change over time we did not find obvious decrease or increase over time. Males and females differed significantly from each other showing nonlinear patterns with females showing a volume decrease until age 20, then increasing again to about their initial volume level. Males increased slightly until age 17, followed by a slight increase.

#### Amygdala

We found no volume change in females but a volume increase in males until about age 16 in males with a following non-linear trajectory. Males also differed significantly from the global trajectory while females did not.

When examining sex differences in each site separately, significant differences between males’ and females’ trajectories were seen for all subcortical structures in Paris and for all subcortical structures except the nucleus accumbens in Dresden (see Figure 3).

**Figure 3.**
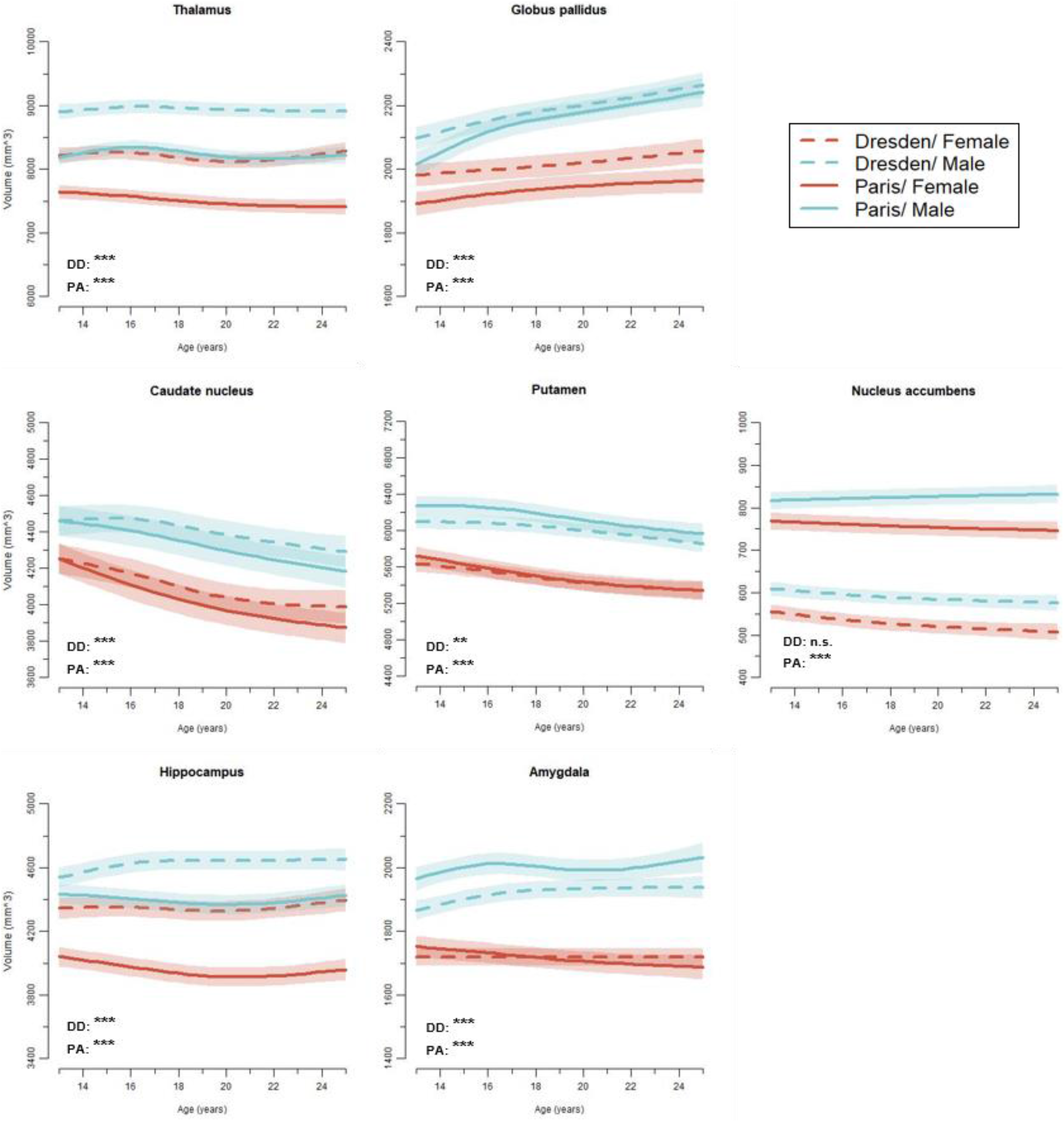
Site differences of developmental trajectories for the subcortical structures. GAMM fitting with 95% confidence intervals plotted separately for females and males in Dresden and Paris. P-values represent sex differences per site; DD = Dresden; PA = Paris; n.s. = not significant; ** p<.01, *** *p* < .001.

### 3.4 Age-constant site differences in volumes

Overall, we found significantly smaller volumes in the Paris site for thalamus, globus pallidus, and hippocampus. Significantly larger volumes in the Paris site were seen for putamen, nucleus accumbens, and amygdala, while the caudate nucleus volume did not differ between the sites.

### 3.5 Site differences in age trajectories

We followed a similar logic for examining the site differences as for the sex differences. For all subcortical structures we found significant differences in age trajectories between the Dresden and Paris site. Examining the sites separately revealed significant volume change in thalamus, globus pallidus, caudate nucleus, and putamen for both sites, significant change only for Dresden in the nucleus accumbens, and no change in both sites for the amygdala.

## 4 Discussion

This study investigated age trajectories and their sex differences across ages 14 to 24 of the volumes of seven subcortical structures in two large samples from Germany and France as part of the IMAGEN project including four waves of data acquisition. This is the first study with a single cohort design to focus on the mid-adolescent to young adult age span providing a continuous data set with 1408 total scans in 503 participants with 36% of participants having three, and 30% of participants having four waves of data acquisition. Our data indicate larger volumes in males compared to females and different age trajectories between the sexes across all subcortical regions. Although the current sample was partly included in the meta-analysis of Dima et al. (2021) we here present all 4 waves from age 14 to 24 processed in the longitudinal stream with FreeSurfer 6.0 (available only for the Dresden and Paris site) versus only the first wave in Dima et al. (2021) was processed using the cross-sectional stream with FreeSurfer 5.3 (available for all IMAGEN sites).

### 4.1 Overall age trajectories across both sexes

We found a decrease in the thalamus from age 16, which was in the range of other studies (15-19 years; Herting et al., 2018; Raznahan et al., 2014; Wierenga, Bos, et al., 2018). Our data show a stabilization thereafter. The slight increase from about age 21 is unexpected and might stem from the few scans at later ages (40 scans after age 23). We found an overall volume increase in the globus pallidus. This is in line with four other studies including a multi-site study (Dennison et al., 2013; Herting et al., 2018; Wierenga, Bos, et al., 2018; Wierenga et al., 2014). Wierenga et al. (2014) had very few scans after age 17, presumably leading to a peak with a following decrease. In contrast, Raznahan et al. (2014), found a decrease. Overall, we found volume decreases in putamen and caudate. This is in line with cross-sectional studies (Brain Development Cooperative Group, 2012; Østby et al., 2009) and a recent mega-analysis (Dima et al., 2021) as well as longitudinal studies (Dennison et al., 2013; Goddings et al., 2014; Tamnes et al., 2013; Wierenga et al., 2014). In contrast, other findings suggest no volume change for the caudate nucleus (Narvacan et al., 2017; Wierenga, Bos, et al., 2018) or an increase for the putamen (Wierenga, Bos, et al., 2018). We demonstrated no volume change in the nucleus accumbens. This is in line with (Dennison et al., 2013), while Wierenga, Bos, et al. (2018) found a decrease. Further, we found a decrease in the hippocampus followed by a slight increase from about age 20. The decreasing pattern is in line with Tamnes et al. (2013) but others found an increase (Dennison et al., 2013; Herting et al., 2018). We demonstrated no volume change in the amygdala which is in line with some previous studies (Dennison et al., 2013; Wierenga, Bos, et al., 2018) but divergent to others (Goddings et al., 2014; Østby et al., 2009; Wierenga et al., 2014). However, findings seem to be driven by sex differences (see e.g. Goddings et al., 2014 and next section on sex differences).

### 4.2 Sex differences

#### 4.2.1 Sex differences in raw volumes

We found larger volumes across subcortical structures for females versus males without allometric scaling. This is in line with previous studies (Herting et al., 2018; Wierenga, Bos, et al., 2018; Wierenga et al., 2014; Wierenga, Sexton, et al., 2018) and the volume differences of 7 to 14 % are in a similar range as found by (Narvacan et al., 2017).

#### 4.2.2 Sex differences in age trajectories

While many previous studies only had few data sets per age group our single cohort study should have the computational strength to precisely model age trajectories during adolescence and detect sex differences. We found significant sex differences in age trajectories for all subcortical ROIs across the two sites. This is in line with Herting et al. (2018) and Raznahan et al. (2014) who also found sex differences in age trajectories for all their subcortical ROIs (Raznahan et al., 2014: striatum, globus pallidus, thalamus). We also found sex differences for each site separately, except for the nucleus accumbens in Dresden. This differs from Herting et al. (2018), who did not find sex differences for all regions when examining their three sites separately. The consistent sex differences for each site might be due to the more homogenous samples and more times points, i.e. power in our study. No sex differences in age trajectories were found in three studies (Narvacan et al., 2017; Wierenga et al., 2014; Wierenga, Sexton, et al., 2018). This is most likely an issue of power (studies covering about 1/ 2, 1/ 6, or 1/ 10 of our total scans) or larger age-ranges (from about primary school until mid-to late twenties). Interestingly, Dennison et al. (2013) had similarly low power (around 1/ 11 of our scans) in a longitudinal design with participants in the adolescent age range and still found sex differences in three subcortical regions. Only Herting et al. (2018) and Raznahan et al. (2014) had similarly powered samples but covered larger age-ranges. Single ROIs will be discussed below.

Overall, descriptively, we observed volume changes for males across several structures until age 16/ 17 while females showed different trajectories in this time span (i.e. stabilization/ decrease). Presumably, females reached the peaks and turning points earlier.

While the thalamus has been shown to change for females or males only (Dennison et al., 2013; Herting et al., 2018), we found a significant change in both sexes. In line with Dennison et al. (2013) and Herting et al. (2018) both sexes first decreased from about age 16 (females) or 17 (males) on. Differing from our study, Dennison et al. (2013) did not cover the age beyond 17. Raznahan et al. (2014) found a rather stable trajectory in males with a slight decrease in females after about age 14. In contrast to our study, Raznahan et al. (2014) had an accelerated design and assessed fewer participants in adolescence while covering earlier ages. Our slight increase from about age 21 is not in line with previous studies and might stem from the few scans after age 23.

For the globus pallidus, females showed a nonlinear increase across the entire age range, while the curve for males increased steeper until age 16 and flattened afterwards. Herting et al. (2018) only found a descriptive albeit non-significant increase for each sex. In contrast, Raznahan et al. (2014) demonstrated a similar slight decrease between the sexes across the adolescent age-range, while Dennison et al. (2013) found an increase in the globus pallidus and no sex differences. Taken together, sex differences in volume age trajectories for the globus pallidus were not convergingly found in the previous literature (also including Narvacan et al., 2017; Wierenga et al., 2014; Wierenga, Sexton, et al., 2018) and our trajectories show an increase in both sexes with just a steeper increase in males.

Since caudate nucleus and putamen form the striatum and show similar trajectories in our data, we discuss them together. We found a decrease for both structures: For females, volume decreased until about age 22 for the the caudate nucleus, respectivley age 20 for the putamen 20, then decrease flattened. Males showed a flatter decrease until about age 17 (putamen 16), followed by a nonlinear decrease. This is in line with Dennison et al. (2013) who found a decrease for both sexes in both structures. No decrease was shown by Herting et al. (2018) for males, but a significant decrease for females in both structures. Raznahan et al. (2014) investigated both structures together as ‘the striatum’ and found an earlier peak for females at age 12 versus 14 for males with a slight decrease thereafter.

In line with Herting et al. (2018) we found a stable trajectory for the nucleus accumbens in males and a decrease in females while other studies (Dennison et al., 2013; Narvacan et al., 2017; Wierenga et al., 2014; Wierenga, Sexton, et al., 2018) did not find sex differences for this region. Taken together, our study together with Herting et al. (2018) is the only one so far to indicate sex differences in the nucleus accumbens.

We discuss amygdala and hippocampus trajectories together since they have similar trajectories. We can only descriptively compare our data with Herting et al. (2018) since they investigated also younger age groups from 8 years on and therefore sex differences in trajectories from age 14 to 24 were not statistically tested. We found no volume change in females but a volume increase in males until about age 16 with a following non-linear trajectory. Males also differed significantly from the global trajectory while females did not. For the amygdala, in line with Herting et al. (2018) who reported stable volumes from age 14 on for females and a slight increase in males, we did not find significant change in females but a nonlinear increase in males. Like Herting et al. (2018) we found significant nonlinear change in the hippocampus for both males and females while our data indicated slight increase from age 20 which was not shown by Herting et al. (2018). Other studies (Dennison et al., 2013; Narvacan et al., 2017; Wierenga, Bos, et al., 2018; Wierenga et al., 2014) did not find sex differences for these regions.

Notably, a recent study investigated sex and puberty effects on structural maturation subcortical structures in a subsample (waves at age 14 and 16) from Dresden and Paris using a voxel-based morphometry and ROI approach (Bézivin-Frere et al., 2020). The authors reported significant differences in volume trajectories with age and puberty in the amygdala-hippocampal complex, caudate, putamen, and thalamus. Together with another study pointing to independent and interactive influences of puberty and age on subcortical development (Goddings et al., 2014) the authors call attention to the importance of pubertal markers when looking at brain development during adolescence, as these processes are likely influenced by differences in sex hormones between females and males. As we only had information on pubertal status for waves one and two and assume pubertal maturation complete afterwards, we did not include pubertal effects in our analyses. Still, differences in subcortical brain development with age and puberty, next to cognitive and social-emotional differences, may add insight into the development of sexual dimorphism in mental disorders during adolescence and later in life (Zahn-Waxler et al., 2008). One mechanism might be the influence of amygdala and hippocampus volume change which differently related to the development of self-reported positive characteristics (e.g. how generous, affectionate, and caring a person is) during adolescence between females and males (Bézivin-Frere et al., 2020).

Reviewing results across all subcortical structures, there is still need for further studies. So far, our study is only the seventh study with three studies spanning only low numbers of 60-147 participants and including only two waves or few participants with three time points (Dennison et al., 2013; Narvacan et al., 2017; Wierenga et al., 2014). All except one study had accelerated designs, so there is a need for larger studies with single cohort designs.

### 4.3 Site effects

Similar to Herting et al. (2018) who also used consistent quality control, longitudinal preprocessing and statistical analyses, we observed site differences and interactions of site by sex depending on the subcortical ROIs. Overall differences in volume between Dresden and Paris were especially visible in thalamus, hippocampus, and nucleus accumbens, and Herting et al. (2018) point to site differences in the thalamus, globus pallidus, caudate, and hippocampus. Interestingly, while Herting et al. (2018) analyzed more heterogeneous sites (differences in number of total scans, age spans, and sex distributions), our two European samples were vastly comparable on sample characteristics such as number of total scans, inter-scan interval, age range, sex distribution and pubertal and socioeconomic status. It is hence unlikely that differences in sample characteristics lead to overall volume differences in Dresden versus Paris. These systematic effects may rather stem from differences in the imaging sequence used on the two sites (higher resolution in Dresden) which may have locally affected the FreeSurfer segmentation algorithm. Similarly, sequence-related differences in subcortical structures have recently been found using CAT-12 (personal communication with J.-L. Martinot). Further, a scanner update was completed during wave three in Paris (Siemens Trio to Prisma) which may have introduced additional bias in results as Medawar et al. (2020) and Plitman et al. (2020) reported very good ICC but also percent volume differences in several cortical and subcortical regions when comparing Siemens Scanners (Trio versus Prisma / Verio versus Skyra). Since many recent longitudinal studies on brain development included several samples with differences in imaging sequence and scanner type (Dennison et al., 2013; Mills et al., 2021; Tamnes et al., 2017; Wierenga, Sexton, et al., 2018), results should be interpreted with caution. Future studies in this field should consider using tools for multi-site harmonization like the recent algorithm longitudinal ComBat shown control type I error rate even better than the unharmonized data with scanner as a covariate approach (Beer et al., 2020).

### 4.4 Limitations

All findings of the present study must be considered in light of the following limitations. Some subcortical structures might suffer from low reliability, i.e. the nucleus accumbens (Mills et al., 2021) putamen or globus pallidus (Wonderlick et al., 2009). The globus pallidus is less distinct from its surrounding white matter as compared to other subcortical regions such as the thalamus or caudate nucleus (Fischl et al., 2002; Wonderlick et al., 2009). Further, the putamen seems to be systematically overestimated in Freesurfer 5.3, which has been improved in the Freesurfer 6.0 (see release notes). This becomes obvious when e.g. comparing putamen volumes at around age 14 in the samples of Herting et al. (2018) of around 6500 mm^3^ for female and 7000 mm^3^ for male with our values (5600 mm^3^ for female 6200 mm^3^ for male).

Concerning GAMM statistics, when global age smooth term is included additionally to group-level smooth term (as done here following Herting et al., 2018) the group-level smooth terms are penalized to not deviate too much from the global smooth terms. This inherently prevents female and male age trajectories from differing too much when fitted in these types of GAMM compared to when fitted separately. Further, concurvity (nonparametric equivalent of multicollinearity) is present when fitting models with separate global and group-level smooth terms as the global term could be approximated by a combination of female and male group-level smooth terms. This might lead to estimation issues in the model and inflated type 1 errors.

Moreover, current results are naturally dependent on power effects: when classes of sexes/ sites are modeled separately the number of data sets is divided in half and significance of effects might be changed (see discussion in Herting et al., 2018). Results further depend on differences in site composition over time as in our study more females than males returned in later waves which might have affected estimation of sex trajectories. Our data also provides more power for the first two waves. Thus, we can be more confident in describing trajectories during mid to late adolescence (until about age 17) while conclusions become less reliable after this time due to attrition of the site (especially after age 23).

While the strength of some previous studies was to show the development in-and-out of adolescence, we focused on the mid-adolescent to young adult age span and provide 1408 total scans within this age span. However, we cannot draw any conclusions about a potential turning point of trajectories before age 14. Additionally, in line with most other studies, the young participants were ethnically homogeneous (i.e. mostly White participants) and stemmed from a well-educated background which restricts generalizability. Future studies could try to recruit participants with a more diverse ethnical and socioeconomic background.

## 5 Conclusions

Our detailed descriptions of regional subcortical trajectories provide further evidence of sex differences in longitudinal adolescent brain development from mid-adolescence to young adulthood. This might eventually add to understanding of the relationship between underlying brain development and different adolescent psychopathology in boys and girls. Additionally, we illuminate pitfalls in large multi-site longitudinal brain imaging studies in an attempt to improve replicability of typical subcortical trajectories over the lifespan.

## Individual contributions

TB, GJB, HF, HG, AH, GS, JLM, and MNS acquired the funding and designed the study.

JHF, NCV and all the other members of the IMAGEN consortium participated in data collection and data management. LLB did the analysis. NCV and LLB drafted the manuscript. NCV, MMH and FS assisted with data analysis and interpretation of findings. All authors critically reviewed the content and approved the final version for publication.

## Acknowledgments

This work received support from the following sources:

the European Union-funded FP6 Integrated Project IMAGEN (Reinforcement-related behaviour in normal brain function and psychopathology) (LSHM-CT-2007-037286), the Horizon 2020 funded ERC Advanced Grant ‘STRATIFY’ (Brain network based stratification of reinforcement-related disorders) (695313), Human Brain Project (HBP SGA 2, 785907, and HBP SGA 3, 945539), the Medical Research Council Grant ‘c-VEDA’ (Consortium on Vulnerability to Externalizing Disorders and Addictions) (MR/N000390/1), the National Institute of Health (NIH) (R01DA049238, A decentralized macro and micro gene-by-environment interaction analysis of substance use behavior and its brain biomarkers), the National Institute for Health Research (NIHR) Biomedical Research Centre at South London and Maudsley NHS Foundation Trust and King’s College London, the Bundesministerium für Bildung und Forschung (BMBF grants 01GS08152; 01EV0711; Forschungsnetz AERIAL 01EE1406A, 01EE1406B; Forschungsnetz IMAC-Mind 01GL1745B), the Deutsche Forschungsgemeinschaft (DFG project numbers 186318919 (FOR 1617), 178833530 (SFB 940), 402170461 (TRR 265), 290210763 (VE 892/2-1), NE 1383/14-1); Faculty of Medicine at the Technische Universität Dresden, MeDDrive Grant; the Medical Research Foundation and Medical Research Council (grants MR/R00465X/1 and MR/S020306/1), the National Institutes of Health (NIH) funded ENIGMA (grants 5U54EB020403-05 and 1R56AG058854-01). Further support was provided by grants from: - the ANR (ANR-12-SAMA-0004, AAPG2019 - GeBra), the Eranet Neuron (AF12-NEUR0008-01 - WM2NA; and ANR-18-NEUR00002-01 - ADORe), the Fondation de France (00081242, 2012-00033703), the Fondation pour la Recherche Médicale (FRM; DPA20140629802; DPP20151033945), the Mission Interministérielle de Lutte-contre-les-Drogues-et-les-Conduites-Addictives (MILDECA), the Assistance-Publique-Hôpitaux-de-Paris and INSERM (interface grant), Paris Sud University IDEX 2012, the Fondation de l’Avenir (grant AP-RM-17-013), the Fédération pour la Recherche sur le Cerveau (FRC Neurodon 2015); the National Institutes of Health, Science Foundation Ireland (16/ERCD/3797), U.S.A. (Axon, Testosterone and Mental Health during Adolescence; RO1 MH085772-01A1), and by NIH Consortium grant U54 EB020403, supported by a cross-NIH alliance that funds Big Data to Knowledge Centres of Excellence.

Resources of The Center for Information Services and High Performance Computing (ZIH) at TU Dresden were used for fast data processing.

We thank Jonas Granzow for help with data processing, quality control and some figures and tables. We further wish to thank Annabell Baake, Leonie Epple, Carolin Fritzsche, and Isabell Theilig for assistance with data management and quality control. Lastly, we thank all participants and their families for their enduring commitment to the study over the years.

