## Supplementary material for "Adolescent to young adult longitudinal development of subcortical volumes in two European sites with four waves"

Table S1. Quality control flow and exclusions. In post processing final data sets and (exclusions) per subcortical structure are listed.

We excluded single subcortical structures when segmented incorrectly. If more than two structures were affected in one image we excluded all structures from quantitative analyses.

**DRESDEN**

| **Pre processing** | **WAVE 1** | **WAVE 2** | **WAVE 3** | **WAVE 4** |
| --- | --- | --- | --- | --- |
| Total MPRAGE scans available | 232 | 218 | 180 | 138 |
| Pre processing QC fail | 30 | 11 | 7 | 8 |
| Available for FS processing | 202 | 207 | 173 | 130 |
| Successfully processed | 202 | 207 | 173 | 130 |
| Post processing QC fail | 1 | 0 | 1 | 0 |
| Total QC fail | 31 | 11 | 8 | 8 |
| Total QC fail percentage | 13.36% | 5.05% | 4.44% | 5.80% |
| Final number of data sets | 201 | 207 | 172 | 130 |

| **Post processing** | **WAVE 1** | **WAVE 2** | **WAVE 3** | **WAVE 4** |
| --- | --- | --- | --- | --- |
| Thalamus | 199 (3) | 207 (0) | 172 (1) | 130 (0) |
| Globus pallidus | 200 (2) | 204 (3) | 172 (1) | 129 (1) |
| Caudate nucleus | 199 (3) | 207 (0) | 172 (1) | 130 (0) |
| Putamen | 200 (2) | 204 (3) | 172 (1) | 129 (1) |
| Nucleus accumbens | 201 (1) | 207 (0) | 172 (1) | 130 (0) |
| Hippocampus | 198 (4) | 206 (1) | 172 (1) | 130 (0) |
| Amygdala | 200 (2) | 206 (1) | 172 (1) | 130 (0) |

**PARIS**

| **Pre processing** | **WAVE 1** | **WAVE 2** | **WAVE 3** | **WAVE 4** |
| --- | --- | --- | --- | --- |
| Total MPRAGE scans available | 257 | 138 | 205 | 123 |
| Pre processing QC fail | 11 | 8 | 4 | 0 |
| Available for FS processing | 246 | 130 | 201 | 123 |
| Successfully processed | 246 | 130 | 201 | 123 |
| Post processing QC fail | 0 | 1 | 1 | 0 |
| Total QC fail | 11 | 9 | 5 | 0 |
| Total QC fail percentage | 4.28% | 6.52% | 2.44% | 0.00% |
| Final number of data sets | 246 | 129 | 200 | 123 |

| **Post processing** | **WAVE 1** | **WAVE 2** | **WAVE 3** | **WAVE 4** |
| --- | --- | --- | --- | --- |
| Thalamus | 246 (0) | 129 (1) | 200 (1) | 123 (0) |
| Globus pallidus | 246 (0) | 129 (1) | 200 (1) | 123 (0) |
| Caudate nucleus | 246 (0) | 129 (1) | 199 (2) | 123 (0) |
| Putamen | 246 (0) | 129 (1) | 200 (1) | 123 (0) |
| Nucleus accumbens | 246 (0) | 129 (1) | 200 (1) | 123 (0) |
| Hippocampus | 245 (1) | 127 (3) | 199 (2) | 123 (0) |
| Amygdala | 245 (1) | 129 (1) | 200 (1) | 123 (0) |

Table S2. Correlations between left and right hemisphere per region of interest for males and females in each wave.

|  | Dresden | |  | Paris | |
| --- | --- | --- | --- | --- | --- |
|  | *r* | *p* |  | *r* | *p* |
| *Thalamus* |  |  |  |  |  |
| Wave 1 – female | 0.83 | *** |  | 0.92 | *** |
| Wave 1 - male | 0.86 | *** |  | 0.91 | *** |
| Wave 2 - female | 0.85 | *** |  | 0.92 | *** |
| Wave 2 - male | 0.89 | *** |  | 0.91 | *** |
| Wave 3- female | 0.83 | *** |  | 0.91 | *** |
| Wave 3 - male | 0.84 | *** |  | 0.86 | *** |
| Wave 4 - female | 0.87 | *** |  | 0.93 | *** |
| Wave 4 - male | 0.89 | *** |  | 0.93 | *** |
| *Globus pallidus* |  |  |  |  |  |
| Wave 1 – female | 0.80 | *** |  | 0.85 | *** |
| Wave 1 - male | 0.72 | *** |  | 0.84 | *** |
| Wave 2 - female | 0.71 | *** |  | 0.87 | *** |
| Wave 2 - male | 0.76 | *** |  | 0.90 | *** |
| Wave 3- female | 0.78 | *** |  | 0.80 | *** |
| Wave 3 - male | 0.78 | *** |  | 0.87 | *** |
| Wave 4 - female | 0.77 | *** |  | 0.83 | *** |
| Wave 4 - male | 0.80 | *** |  | 0.86 | *** |
| *Caudate nucleus* |  |  |  |  |  |
| Wave 1 – female | 0.94 | *** |  | 0.95 | *** |
| Wave 1 - male | 0.92 | *** |  | 0.95 | *** |
| Wave 2 - female | 0.94 | *** |  | 0.95 | *** |
| Wave 2 - male | 0.92 | *** |  | 0.96 | *** |
| Wave 3- female | 0.94 | *** |  | 0.94 | *** |
| Wave 3 - male | 0.94 | *** |  | 0.95 | *** |
| Wave 4 - female | 0.94 | *** |  | 0.94 | *** |
| Wave 4 - male | 0.94 | *** |  | 0.96 | *** |
| *Putamen* |  |  |  |  |  |
| Wave 1 – female | 0.89 | *** |  | 0.96 | *** |
| Wave 1 - male | 0.91 | *** |  | 0.97 | *** |
| Wave 2 - female | 0.91 | *** |  | 0.96 | *** |
| Wave 2 - male | 0.87 | *** |  | 0.97 | *** |
| Wave 3- female | 0.88 | *** |  | 0.93 | *** |
| Wave 3 - male | 0.92 | *** |  | 0.96 | *** |
| Wave 4 - female | 0.91 | *** |  | 0.94 | *** |
| Wave 4 - male | 0.90 | *** |  | 0.97 | *** |
| *Nucleus accumbens* |  |  |  |  |  |
| Wave 1 – female | 0.67 | *** |  | 0.67 | *** |
| Wave 1 - male | 0.46 | *** |  | 0.73 | *** |
| Wave 2 - female | 0.68 | *** |  | 0.66 | *** |
| Wave 2 - male | 0.46 | *** |  | 0.74 | *** |
| Wave 3- female | 0.72 | *** |  | 0.65 | *** |
| Wave 3 - male | 0.53 | *** |  | 0.68 | *** |
| Wave 4 - female | 0.69 | *** |  | 0.72 | *** |
| Wave 4 - male | 0.51 | *** |  | 0.79 | *** |
| *Hippocampus* | 0.80 | *** |  | 0.86 | *** |
| Wave 1 – female | 0.83 | *** |  | 0.84 | *** |
| Wave 1 - male | 0.76 | *** |  | 0.90 | *** |
| Wave 2 - female | 0.83 | *** |  | 0.81 | *** |
| Wave 2 - male | 0.80 | *** |  | 0.85 | *** |
| Wave 3- female | 0.78 | *** |  | 0.86 | *** |
| Wave 3 - male | 0.81 | *** |  | 0.83 | *** |
| Wave 4 - female | 0.80 | *** |  | 0.85 | *** |
| Wave 4 - male | 0.80 | *** |  | 0.86 | *** |
| *Amygdala* |  |  |  |  |  |
| Wave 1 – female | 0.72 | *** |  | 0.69 | *** |
| Wave 1 - male | 0.75 | *** |  | 0.72 | *** |
| Wave 2 - female | 0.75 | *** |  | 0.76 | *** |
| Wave 2 - male | 0.77 | *** |  | 0.73 | *** |
| Wave 3- female | 0.77 | *** |  | 0.68 | *** |
| Wave 3 - male | 0.77 | *** |  | 0.71 | *** |
| Wave 4 - female | 0.74 | *** |  | 0.72 | *** |
| Wave 4 - male | 0.79 | *** |  | 0.60 | *** |

*** *P-value* <.001.

Table S3. AIC values for each structure for models 1 - 3.

| Structure | Model 1 | Model 2 | Model 3 |
| --- | --- | --- | --- |
| Thalamus | 19719.76 | 19564.18 | 19541.10 |
| Globus Pallidus | 16524.41 | 16427.87 | 16278.20 |
| Caudate Nucleus | 18034.79 | 17984.28 | 17851.52 |
| Putamen | 18601.97 | 18459.73 | 18412.57 |
| Nucleus Accumbens | 14950.59 | 14906.73 | 14884.50 |
| Hippocampus | 17797.75 | 17677.13 | 17631.97 |
| Amygdala | 16585.26 | 16391.31 | 16325.84 |
